# Inverse signal importance in real exposome: How do biological systems dynamically prioritize multiple environmental signals?

**DOI:** 10.1101/2025.03.31.646257

**Authors:** Thoma Itoh, Yohei Kondo, Tomoya Nakayama, Ai Shinomiya, Kazuhiro Aoki, Takashi Yoshimura, Honda Naoki

## Abstract

Living organisms integrate multiple signals from their exposome, the totality of environmental influences experienced throughout life, to adapt to complex, non-stationary environments. While organisms are thought to flexibly prioritize relevant signals depending on context, its regulatory mechanisms remain largely unknown. Laboratory studies with precisely controlled conditions fail to capture this adaptability by isolating organisms from the complex exposome. Here, we developed a machine learning framework, Inverse Signal Importance (ISI), to infer how organisms prioritize external signals from time-series data of environmental factors and physiological responses. We applied ISI to analyze gonadal development in medaka fish under natural outdoor conditions, tracking gonadosomatic index alongside environmental signals including water temperature, day length, and solar radiation over two years. Our analysis revealed that signal importance levels exhibit complex dynamics distinct from simple environmental periodicity and correlates significantly with specific gene expression patterns. Notably, genes associated with temperature-related signal importance display differential expression between outdoor and controlled laboratory conditions, suggesting their role in environmental adaptation. These findings indicate that ISI effectively captures latent physiological dynamics in adaptation of exposome. By decomposing biological responses into deterministic and adaptive components, ISI provides a novel approach to uncover mechanisms of organismal adaptation in natural environments.

## Introduction

Living organisms make decisions by transforming and integrating multiple streams of information in an uncertain world. The overwhelming complexity has been captured through the notion of *exposome*, which encompasses the totality of environmental influences and associated physiological responses throughout the lifespan [1, 2].

Across scales, the biological responses under the complex environment emerge from selective use of diverse signals: animals determines the behavior by flexibly attending to sensory modalities such as auditory, visual, and olfactory signals depending on context and experience [3, 4, 5]. Within the organism, metabolic systems flexibly allocate resources towards processes such as organ growth, reproduction and immune response to infection depending on the environment [6, 7, 8]. Even single cells integrate not only ligand concentrations but also their spatial and temporal patterns and molecular composition [9, 10]. These observations motivate a central question: how do organisms flexibly integrate and/or select environmental signals in an ever-changing world [11, 12, 13, 14]?

A natural starting point for formalizing this question is information processing of time-varying signals. In many systems, an input signal is not used instantaneously but is temporally integrated through a response function [15, 16, 17], which can be represented as a convolution (Fig. 1**a**). With multiple input signals, a standard extension is to filter each modality and then integrate them by a weighted sum (Fig. 1**b**). Such models have been widely used to capture sensory integration, neural responses, and molecular decoding of complex stimuli [18, 19]. However, a key assumption is often implicit that the integration weights are constant, meaning that the system’s sensitivity to each signal modality does not change over time. This assumption is convenient in controlled laboratory settings but becomes problematic in real environments, where non-stationary inputs (*e*.*g*., seasonal temperature and photoperiod changes, rainfall, predation risk, and other factors) continually reshape the informational context [20]. In such settings, organisms are expected to update the relative importance assigned to different input signals (Fig. 1**c**).

**Figure 1.**
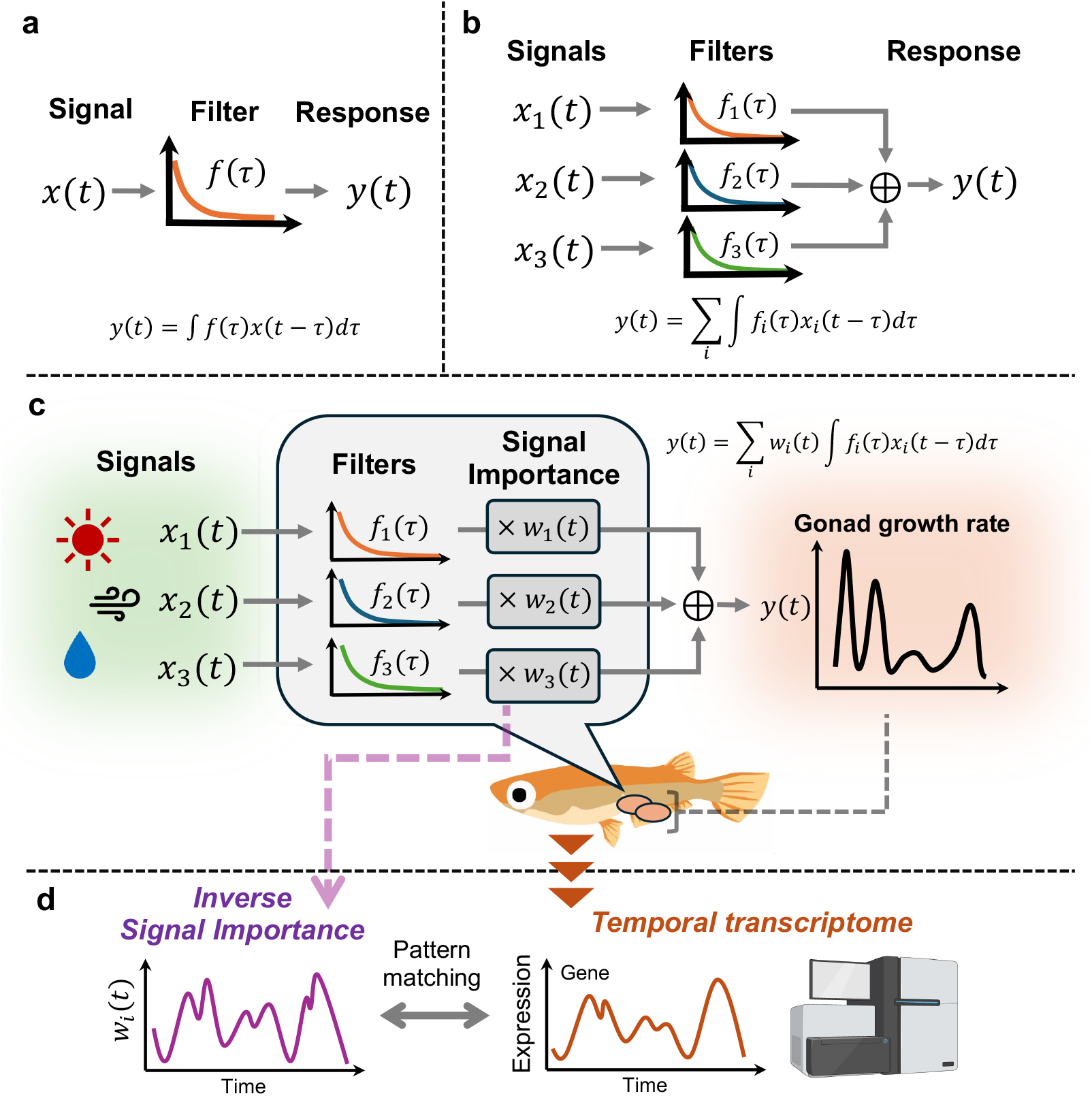
Conceptual framework for the analysis of biological response for environment. **a** Single signal model. Input signal *x*(*t*) is processed through a filter *f*(*τ*) to generate a response *y*(*t*), described by the convolution integral *y*(*t*) = ∫ *f*(*τ*)*x*(*t* − *τ*)*dτ*. The response at the current time *y*(*t*) is calculated as a weighted sum of past input signals *x*(*t* − *τ*), where the kernel *f*(*τ*) determines the weight based on the time lag *τ*. This mathematical framework captures the cumulative and delayed nature of biological responses to environmental stimuli. **b** Multiple signal model. A more complex model where multiple signals *x*_*i*_ are processed by individual filters *f*_*i*_(*τ*). The final response *y*(*t*) is the summation of these filtered components. **c** Multiple signal model with signal importance. Signal Importance modulates the weights of each signal modalities to represent time-dependent sensitivity to specific environmental signals. This framework can capture the flexibility of biological outputs such as gonad size of medaka under the non-stationary environment. **d** Inverse signal importance and identification of environment-responsive genes. Through the “Inverse Signal Importance” method, the underlying dynamical state *w*_*i*_(*t*) are inferred (see methods for details). Pattern matching between these inferred states and specific gene expression profiles enables the identification of genes that are likely responsive to environmental adaptation. Created with BioRender.com.

Our recent work demonstrated both the strength and limitations of the time-invariant framework. We used field data to infer how multiple environmental cues are temporally processed and integrated to shape a physiological output. Analyzing robust annual changes in gonad size in medaka fish (*Oryzias latipes*) under uncontrolled outdoor conditions, we extracted environmental dependencies of the gonad growth by estimating how photoperiod, temperature, and other cues are processed through temporal integration [21]. However, both predictive performance and interpretability were limited by the time-invariant formulation: the model effectively assumed that the cue-integration from environmental inputs to gonadal dynamics remains fixed across the year. This assumption is unlikely to hold in medaka’s natural conditions, because photoperiod and temperature are known to be processed differently across seasons and physiological states in their natural life history [21, 22]. More broadly, it highlights a general gap between classical multi-input integration models and the flexible, context-dependent signal selection that real organisms exhibit.

Here we introduce Inverse Signal Importance (ISI), a framework that explicitly models and infers time-varying signal importance from observational time series (Fig. 1**c**). ISI augments conventional multi-input integration by replacing constant weights with a latent dynamical variable ***w***(***t***) that represents the instantaneous importance of each signal modality. Because ***w***(***t***) is not directly observable and emerges from complex interplay between physiological system and a non-stationary environment, inferring it from data constitutes an inverse problem. ISI addresses this challenge with a hybrid approach that combines (i) time-invariant temporal filters ***f*** and (ii) time-varying weights ***w***(***t***) reconstructed via state-space inference (Kalman filtering) using recorded inputs and observed physiological outputs.

Building on this formulation, we organize our study around two steps (Fig. 1 **c**,**d**). First, we decode signal importance by reconstructing the constant filter ***f*** and time-variable ***w***(***t***) (signal importance) in a data-driven manner from field environmental time series and a physiological output (Fig. 1**c**). We interpret the inferred ***w***(***t***) as dynamic prioritization, quantifying when and how the organism re-weights environmental signals across seasons and contexts. Second, we biologically ground this latent prioritization by linking ***w***(***t***) to time-series transcriptome measurements, thereby identifying the candidate environment-responsive genes whose expression dynamics accordance with the inferred signal importance ***w***(***t***) (Fig. 1**d**). Applying this pipeline to circannual gonad-size dynamics in medaka under outdoor conditions, ISI provides a general route from non-stationary exposome data to prioritization for signal modalities and their candidate molecular implementations.

## Results

This study aims to quantify the importance of signals by focusing on the gonadal development of medaka fish through the inverse signal importance (ISI) method. Previously, we have characterized living organisms through static models that quantify the relationship between physiological inputs and outputs [21]; the ISI method extends the framework by introducing internal states for signal importance that weight outputs of mechanistic models. The extension enables us to infer how living systems change which signals to focus on.

### Gonad size in medaka under natural outdoor conditions

We previously measured the gonad size of medaka fish every two weeks over two years (October 2015 to October 2017) under natural outdoor conditions in Okazaki, Japan (Fig. 2**a**) [21]. The time evolution of the gonadosomatic index (GSI), defined as gonad weight as a percentage of body weight, exhibited a noticeable annual rhythm (Fig. 2**a**, bottom panel). In the current study, we trained a predictive model, illustrated in Figure 2**b**, for the differentiation of GSI (dGSI) as the output variable (Fig. 2**c**), while the environmental data (solar radiation, water temperature, and day length) and past GSI values are used as input variables (Fig. 2**a**). Notably, the inclusion of the past GSI value as an input for prediction aligns with widely established practices in machine learning.

**Figure 2.**
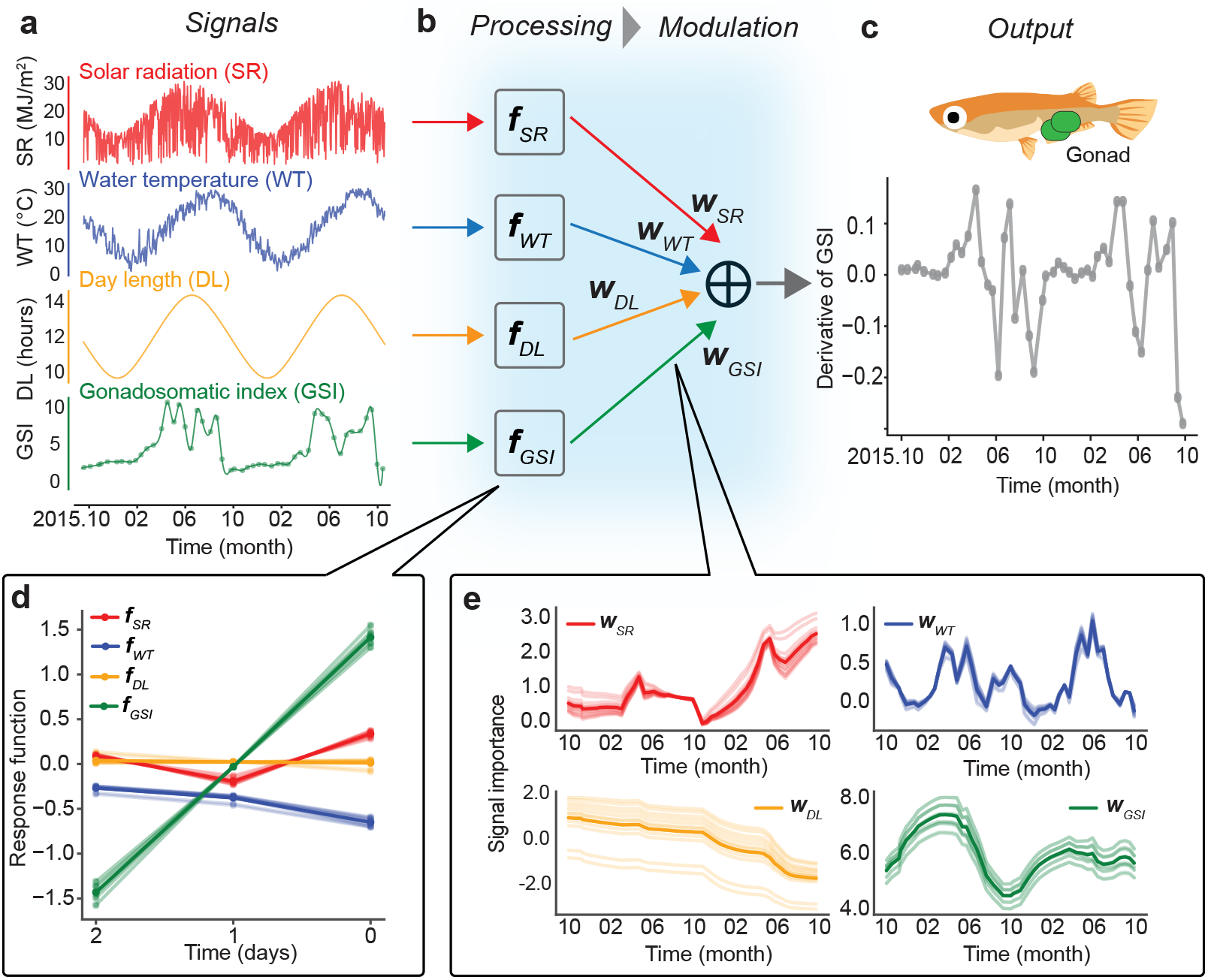
**a**-**c** Framework overview. Environmental data and gonad somatic index (**a**) are processed through a linear filter ***f***, followed by the modulation of signal importance by multiplication with ***w*** (**b**). Each signal has corresponding ***f*** and ***w*** (SR: solar radiation, WT: water temperature, DL: day length, GSI: gonad somatic index), which are integrated to predict the derivative of GSI at the immediate future time point (**c**). Panel **a** shows the temporal changes in environmental data and gonad somatic index (GSI) as input variables from October 2015 to October 2017, while panel **c** shows the temporal changes in the derivative of GSI as output variables over the same period. **d, e** The estimated variables: the optimal response linear filters (**d**) and the optimal signal importance weights (**e**). Thin pale lines correspond to models with comparable performance; thick solid lines are the averages. Note the distinct temporal scales between the response function (days) and the signal importance (month).

### Signal importance on gonad size regulation of medaka

In our ISI framework (Fig. 2), each environmental signal is processed separately according to corresponding processing rule, which is defined by time-invariant linear filter ***f*** . Then the processed signals are weighted by signal importance ***w***(*t*) (See Methods and Materials for details). We developed a machine learning method to infer ***f*** and ***w***(*t*), and applied it to the time-series data of medaka physiology and environmental signals (see Supplementary Note and Supplementary Figs. 1 and 2). Then we successfully identified how medaka process the environmental signals (***f***) and prioritize them (***w***) to regulate gonadal size. We validated the estimation by confirming the robustness against variation of hyper-parameters (Fig. 2**d**,**e**; pale lines). The profile of the processing rule is consistent with our previous study [21] showing flat patterns except for the ***f***_*GSI*_ (Fig. 2**d**). The ***f***_*GSI*_ exhibits the characteristics of a differentiator, negatively referencing value of 2 days ago, but positively referencing values of 0 days ago, suggesting that the change of gonad somatic size is highly affected by the past trajectory of gonad somatic size. The optimized trajectories of ***w***_*SR*_ and ***w***_*WT*_ show deviations from the near-perfect periodicity of environmental signals, (Fig. 2**e**), suggesting the complexity of information processing. On the other hand, ***w***_*DL*_ shows a monotonic decrease, and ***w***_*GSI*_ shows a simple sinusoidal trajectory (Fig. 2**e**). Surprisingly, this estimation revealed that the periodic pattern of the temporal derivative of GSI emerges through asynchronous dynamics of ***w***. This indicates that the regulatory system continuously recalibrates its signal integration in response to environmental fluctuation, demonstrating a homeostatic capacity for maintaining periodic rhythm of gonadal development under variable conditions.

### Concordance between gene expression patterns and signal importance

Previously we collected monthly transcriptome for over a two-year period, which was simultaneously sampled with the GSI and environmental data [21]. Based on our hypothesis that the estimated ***w***_*i*_ (*i*=*SR, WT, DL, GSI*) values reflect environmental adaptation, we expected to find genes exhibiting similar dynamics with ***w***_*i*_. Hence, we compared the time-series transcriptome with each ***w***_*i*_ except for the ***w***_*DL*_ which monotonically decreased throughout the entire period. As a result, we found several genes showing high concordance with the ***w*** (Fig. 3**a**). The similarity between gene expression pattern and ***w*** was quantified with the correlation coefficient. The correlation coefficients across all genes displayed a trapezoid distribution (Fig. 3**b**).

**Figure 3.**
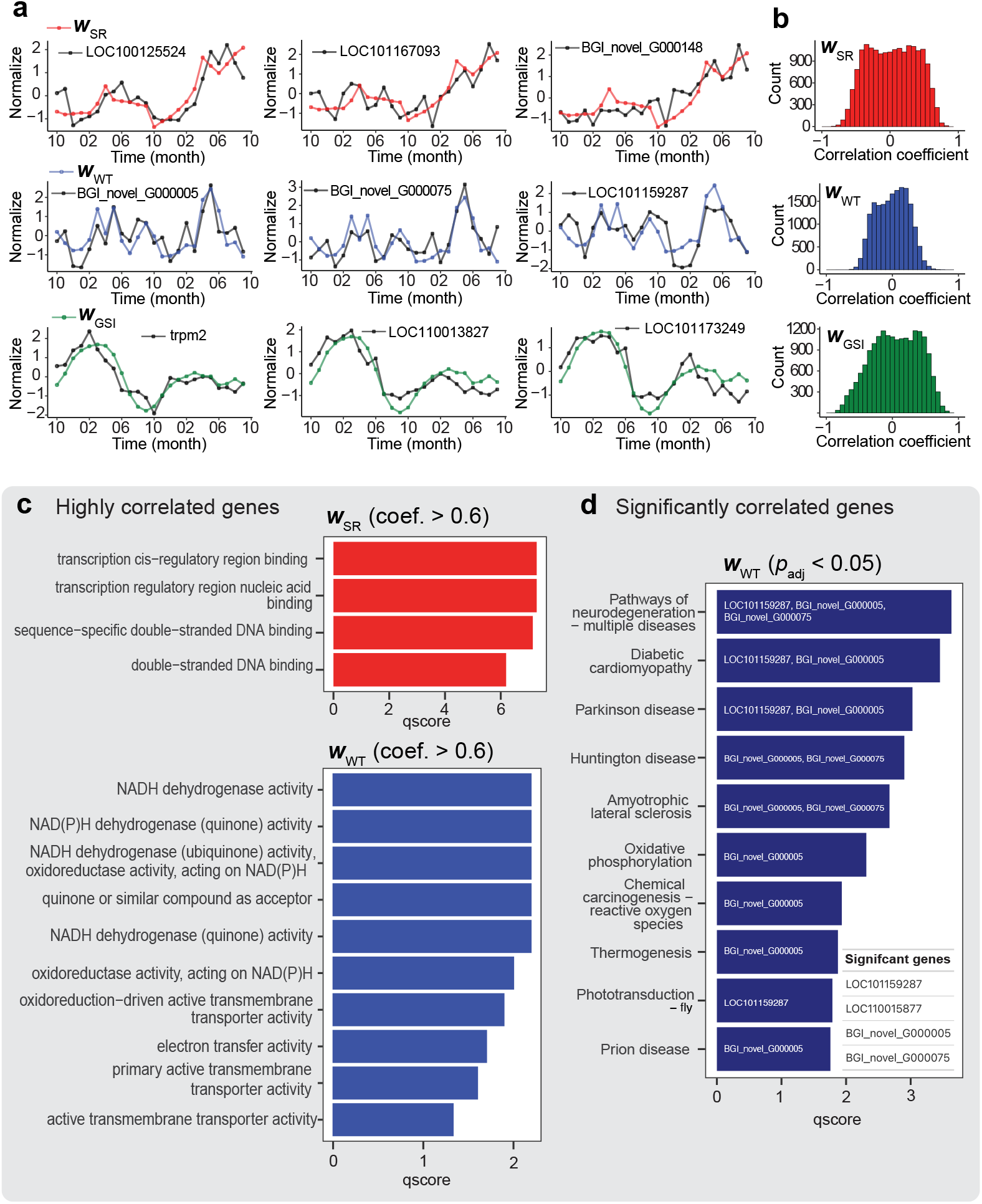
**a** The genes synchronized with the signal importance. The expression level (black) and signal importance (***w***_*SR*_: red, ***w***_*WT*_: blue, ***w***_*GSI*_: green) are normalized. The expression levels represent the average of two technical replicates. **b** Distribution of correlation coefficient between ***w*** and all genes (The genes which have missing value are excluded). **c**,**d** Enrichment analysis. The horizontal axis shows the significance of the enrichment with qscore (-log10p) and the vertical axis shows the enriched function (qscore > 1). **c** GO enrichment analysis (Molecular function) for genes highly correlated (coef. > 0.6) with ***w***_*SR*_ (red) and ***w***_*WT*_(blue). **d** KO enrichment analysis was performed on highly correlated genes that showed statistical significance (p_adj_ < 0.05; Bonferroni correction). Among the tested genes, only those correlated with ***w***_*WT*_ passed the statistical test. The list of significantly correlated genes is listed in the lower right table, and the gene names corresponding to each function are annotated on the bar.

To investigate the functional tendency of the highly correlated genes, we extracted the genes which show correlation coefficients higher than 0.6 and then applied them for the enrichment analysis (clusterProfiler in R package) [23]. The genes correlated with ***w***_*SR*_ are enriched in damage response categories (e.g., double-strand binding) and the genes correlated with ***w***_*WT*_ are enriched in energy generation-related terms (e.g., NADH activity, cellular respiration) in GO enrichment analysis (FDR 0.1; molecular function) (Fig. 3**c**). All GO enrichment results including biological process are shown in the supplementary figure 3.

To further narrow down the candidate genes, we excluded genes that showed spurious high correlations due to random chance. To this end, we computed correlation coefficients between each gene expression pattern and randomized trajectories of the signal importance, which allowed us to assess the significance of the observed correlation to randomized ones (Supplementary Fig. 4). Genes with *p* < 0.05 after Bonferroni correction were selected. The correlation coefficients, along with both raw and adjusted *p*-values for the highly correlated genes, are provided in Supplementary Table 1.

**Figure 4.**
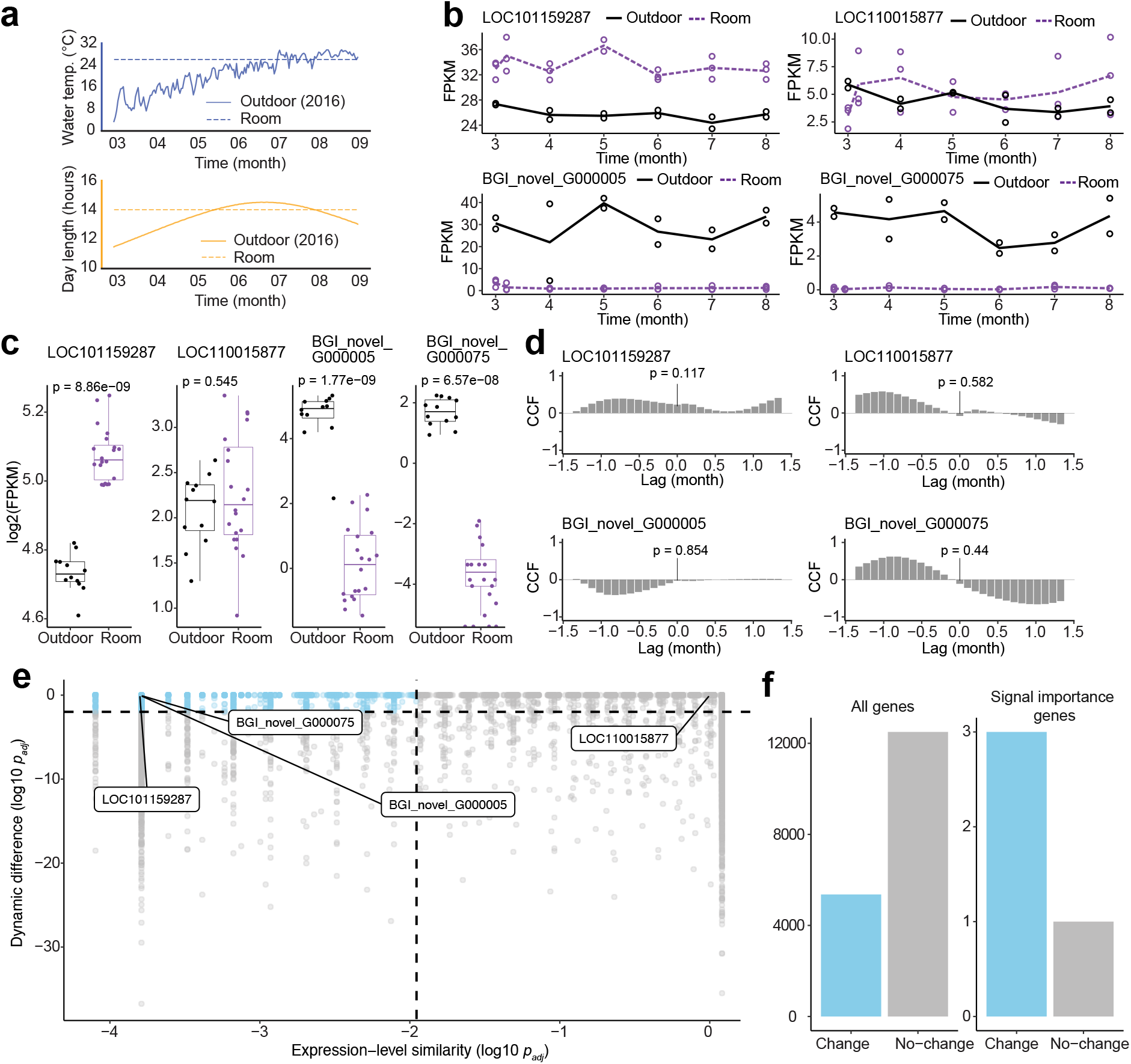
Expression dynamics of signal importance genes differ between outdoor and room conditions. **a** Environmental signals in outdoor (solid line) and room (dashed line) conditions. The upper panel represents water temperature, and the lower panel shows day length. The room condition was maintained at a constant 14-hour day length and 26°C water temperature. **b** Temporal patterns of expression (FPKM) in the outdoor (black solid line) and room (purple dashed line) on the signal importance genes that are significantly correlated with water temperature. Dots show RNA-seq samples, and lines show averages in outdoor (black: 2 samples) and room (purple: 3 samples). **c** Expression level distributions across time series and samples. Box plots compare expression levels between outdoor (left) and room (right) conditions. Wilcoxon signed-rank test p-values are shown above. **d** Cross correlation between the outdoor and room expression levels. Spearman rank correlation p-value at lag 0 indicated. **e** Comparison between expression level similarity and temporal pattern difference across conditions. X-axis shows Wilcoxon test p-values indicating expression level similarity between conditions (as in **c**); Y-axis shows Spearman test p-values reflecting differences in temporal expression patterns between conditions (as in **d**). Points represent genes. Genes located in the upper left quadrant (highlighted in blue) are considered to exhibit different expression dynamics. **f** The ratio of the genes located in the upper left region of panel **e**. Comparison between all genes (left) and signal importance genes (right). Fisher’s exact test suggests a marginal difference (p=0.058).

As a result, four genes were selected as significantly correlated with ***w***_*WT*_ (hereafter referred to as signal importance genes). Note that any genes did not show the significant correlation with ***w***_*SR*_ and ***w***_*GSI*_. Signal importance genes include two known genes (LOC101159287, LOC110015877) and two novel transcripts (BGI_novel_G000005, BGI_novel_G000075). Notably, GO enrichment analysis revealed these genes to be associated with energy generation (Supplementary Fig. 3**f, g**), while KO enrichment highlighted their involvement in thermogenesis (Fig. 3**d**). The enrichment analyses indicate synchronization between thermoregulation-related genes and the signal importance of water temperature (***w***_*WT*_), suggesting seasonal physiological adaptation in medaka. These findings support that signal importance ***w*** reflects change of the internal state of the living system.

### Signal importance genes are involved in environmental adaptation

We hypothesized that the signal importance genes, which are significantly correlated with the seasonal change of signal importance, play key roles in environmental adaptation. To test the hypothesis, we compared their expression dynamics under the non-stationary environment (outdoor condition) and the stationary environment (room condition). The expression data in the stationary environment were collected over 6 months under controlled room conditions (14 h daylight and 26 °C water temperature; Fig. 4**a**) in our previous study [21]. Although the GSI pattern shows the similar bimodality in the room and outdoor condition (Nakayama et al., 2023) [20], the expression patterns show clear separation between the room and outdoor conditions in most signal importance genes for water temperature (Fig. 4**b**). In the room condition, the signal importance genes except LOC110015877 show distinct expression levels (Fig. 4**c**). The correlation of expression patterns between room and outdoor conditions are low and non-significant, showing differences in expression patterns (Fig. 4**d**). The three genes (LOC101159287, BGI_novel_G000005, and BGI_novel_G000075) show distinct expression patterns – both in their absolute levels and temporal changes – suggesting that these genes operate in distinct modes under stationary versus non-stationary environments (Fig. 4**d**).

To assess how the signal importance genes differ in their expression patterns, we analyzed all genes using the same procedure applied to signal importance genes in Figures 4**c** and **d**. We quantified both the similarity in absolute expression levels between conditions (Fig. 4**e**, x-axis) and the differences in expression trajectories (Fig. 4**e**, y-axis). Genes in the upper left region of Figure 4**e** (Wilcoxon rank-sum test, p < 0.01; Spearman correlation, p > 0.01; Bonferroni-corrected) exhibit significant differences in both expression levels and temporal changes between conditions. Conversely, genes in the lower right region of Figure 4**e** exhibit similar dynamics under both conditions, including the ‘circannual genes’ that display free-running oscillation regardless of condition (Supplementary Figure 5), as previously identified (Nakayama et al., 2023) [21]. Among genes significantly correlated with signal importance, 75% (3/4) are in the upper left region, compared to only 26% (4,708/13,153) of all genes (Fig. 4**f**). The difference between the ratios is marginally significant (Fisher’s Exact Test, p = 0.058), suggesting that the latent dynamics deduced from our ISI method captures the dynamics of genes involved in the environment adaptation.

### Association with the sex hormone genes

Because our analysis derived signal importances as modulators of gonadal growth, we investigated whether these weights covary with the temporal expression trajectories of sex-hormone genes. We assembled a reference set of 15 sex-hormone transcripts, including estrogen-receptor isoforms, aromatase, androgen-synthesis enzymes, and other steroidogenic factors [24, 25, 26, 27, 28, 29, 30, 31, 32, 33, 34] (Supplementary Table 2). We then compared their temporal expression trajectories with the time-series of the inferred weights (***w***_*SR*_, ***w***_*WT*_, ***w***_*GSI*_) and environmental signals (SR, WT, GSI, dGSI). This cross-comparison revealed covariation between the environmental signals and sex-hormone transcripts (Supplementary Fig. S6). For example, water temperature closely correlate with the expression of *hsd17b3* and *hsd17b4* (*r* > 0.85), and solar radiation show strong concordance with *cyp19b* (aromatase; r > 0.68) as well as with *pgr* and *hsd17b3*, which are known as genes central to ovulation and testosterone synthesis respectively [33, 29] (r > 0.57). In contrast, We found no gene whose expression uniquely tracked the temporal dynamics of ***w***_*SR*_, ***w***_*WT*_, or ***w***_*GSI*_. As an exceptional case, while ***w***_*SR*_ roughly correlate with the raw SR profile and aligned with the same transcripts (*cyp19b, pgr* and *hsd17b3*; r > 0.68), this alignment is not signal importance–specific. Taken together, these findings suggest that the signal-importance weights do not mirror the direct action of sex-hormone genes. Rather, they reflect an indirect tuning of gonadal growth mediated through environmental adaptations, as evidenced by the strong covariation between signal importance and metabolic or immune-related transcripts in Figure 3.

## Discussion

Our study introduces the inverse signal importance (ISI), a novel framework to infer how living organisms dynamically prioritize multi-modal environmental signals. We applied the method to the gonadal development of medaka fish. The inferred signal importances exhibit complex trajectories that deviate from simple annual periodicity, and these trajectories show strong correlations with expression patterns of several genes. In particular, the signal importance for water temperature (***w***_*WT*_) showed a significant correlation with four genes, three of which display distinct expression dynamics between outdoor and room conditions, indicating their potential role in environmental adaptation. In contrast, although the signal importances were inferred as modulators of gonadal growth, none exhibited covariation with sex-hormone transcripts, indicating that signal importances reflect an indirect mode of growth regulation mediated through pathways of environmental adaptation rather than direct hormonal control.

In our previous work, we characterized circannual genes that maintain oscillatory patterns across environments, providing novel insights into the robust regulation of seasonal rhythms. In contrast, ISI identifies genes whose expression dynamically changes in a physiologically meaningful pattern (signal importance, ***w*** ), derived from the relationship between input signals and output responses. For example, genes significantly correlate with the signal importance of water temperature (***w***_*WT*_) exhibited adaptive transcriptional responses that differed between outdoor and laboratory conditions (Fig. 4**e**,**f**). These findings underscore the utility of ISI in disentangling context-dependent adaptation mechanisms from broader patterns of environmental responsiveness.

However, not all inferred signal importances showed gene associations. For signal importances other than water temperature, none shows a significant correlation with any genes. This raises the possibility that these signals may came from confounding factors. For example, genes highly (though not significantly) correlated with the solar radiation importance (***w***_*SR*_) exhibit monotonic increases during the second year despite relatively stable inputs (SR), and was weakly associated with genes enriched for immune-related terms (Fig. 3). These patterns may indicate unmeasured environmental change such as shifts in water quality or microbial activity which leads to compensatory physiological responses. These observations underscore the need for further investigation to interpret the nature of the inferred signal importances.

Living systems demonstrate robustness across multiple scales, from gene expression [35] to annual rhythms [21]. Hence, the superficial traits (i.e., rhythms or morphology) often appear unchanged even under perturbations due to internal compensatory mechanisms. For example, the circadian rhythm of *Arabidopsis thaliana* maintains robust oscillations against temperature fluctuations through the temperature-responsive genes, which counterbalance the effects of temperature changes on biochemistry [36]. The ISI framework offers a valuable lens to understand such internal mechanisms by separately modeling signal processing systems and its modulators. Additionally, beyond ecological relevance, the ISI framework may offer pharmacological insights by detecting latent physiological states shaped by environmental exposures. For example, an increase in ***w***_*SR*_ that deviates from the stable seasonal oscillation of solar radiation while aligning with immune-related gene expression, suggests an internal stress response. Such signal-importance profiles may serve as early indicators of environmentally induced physiological dysregulation. As environmental factors are increasingly recognized as modulators of drug metabolism and efficacy, ISI provides a novel approach for linking external environmental signals to latent physiological states (signal importance), which may inform new pharmacological strategies depending on the exposome context.

In summary, ISI decomposes complex environmental time series into deterministic and adaptive components. While previous studies may have dismissed residual variations as noise, our framework reinterprets these variations as reflections of internal adaptation processes. The findings from medaka fish gonadal development exemplify how organisms adjust internal regulatory systems in response to environmental changes. Therefore, the ISI framework paves the way to reveal organismal adaptation to the natural environment.

## Methods

### A. Data processing

#### Meteorological and GSI data

We used the preprocessed meteorological and GSI data from the previous study (Nakayama et al., 2023). The data set was collected at the National Institute for Basic Biology in Okazaki from October 1, 2015, to October 15, 2017, with sampling conducted twice per month. In each sampling, the GSI of ten female medaka was recorded. For the calculation of derivative of GSI 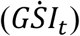, the time series of GSI are first averaged over medaka individuals. Subsequently, the derivative at each sample point was computed through the linear fitting (Supplementary Fig. 1).

#### Data splitting

To prevent overfitting, we divided the GSI data from 10 female specimens into three subsets: Data 1 (n = 3), Data 2 (n = 3), and Data 3 (n = 4) (Supplementary Fig. 1). During hyperparameter tuning, we employed cross validation, i.e., using two subsets for training and the remaining one for testing (Supplementary Fig. 2 b). Once the optimal hyperparameters were determined, the entire dataset was utilized to train the final model and estimate the parameters.

#### Transcriptome data

Low-quality sequencing reads were filtered out. Genes with missing data on any time point were excluded from downstream temporal analyses.

### B. Mathematical model

#### Inverse Signal Importance

The ISI method is based on the following model:

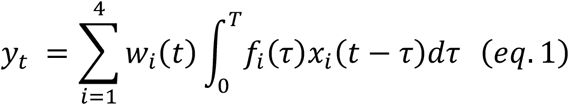

where *y*_*t*_ is the temporal differentiation of GSI at time = *t* 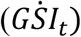 in this study. The input signal *x*_*i*_(*t*) indicates solar radiation (SR, *x*_1_ ≡ *x*_*SR*_), water temperature (WT, *x*_2_ ≡ *x*_*WT*_), day length (DL, *x*_3_ ≡ *x*_*DL*_), and Gonadosomatic index (GSI, *x*_4_ ≡ *x*_*GSI*_), at time = *t*. We include the past GSI value as input to predict output 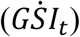 as widely utilized in machine learning. *f*_*i*_(*τ*) is a time-invariant regression parameter for the input *i* and delay *τ*, and *w*_*i*_ (*t*) is a time-varying coefficients for the input *i* at time *t* . The convolution of past signal information, 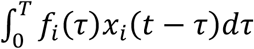, is known as an ordinary response function with kernel size *T*. The response function is further modified by the factor *w*_*i*_(*t*) which vary over time to describe seasonal modulation.

#### Parameter inference framework: hard EM algorithm

For parameter inference, let us consider the time discretization of eq. 1 as

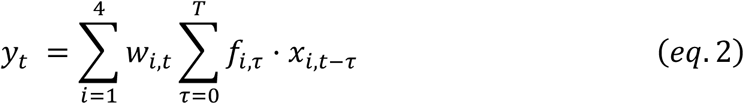

where *x*_*i,t*_, *f*_*i,t*_, and *w*_*i,t*_ represent *x*_*i*_(*t*), *f*_*i*_ (*t*), and *w*_*i*_ (*t*). For brevity, we denote by ***x*** the collection of all *x*_*i,t*_ variables for all *i* and *t*. The same notation applies to ***f*** and ***w***.

The parameters ***f*** and ***w*** are inferred to maximize the following log-likelihood:

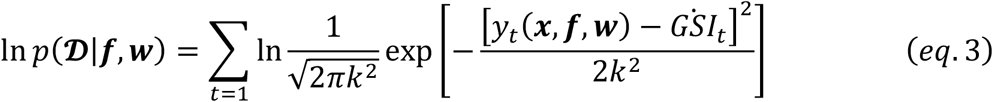

where *y*_*t*_(***x, f, w***) is a predicted *GSI*_*t*_ from our model (eq. 2), and 𝒟 is a dataset including ***x*** and 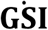.

We adopted an alternating optimization method for the log-likelihood, which is inspired by the EM algorithm (For complete derivation, see Supplementary Note 1) [37].

#### E step: Optimization of w

Signal importance ***w*** is inferred with the following state-space model under fixed ***f***^*^:

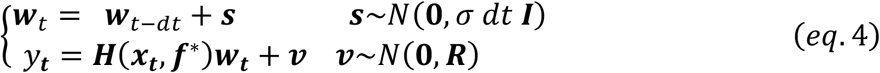

where ***H***_***t***_(***x***_***t***_, ***f***^*^) is the set of processed signals through a linear filter ***f***^*^, ***s*** is process noise, and ***v*** is observation noise (Supplementary Note2). Here we used ***x***_***t***_ to represent [*x*_1,*t*_, *x*_2,*t*_, *x*_3,*t*_, *x*_4,*t*_]. Based on *eq*. 4, the optimal ***w*** is inferred by the Kalman filter. The variance of ***v*** is estimated from the data (Supplementary Note 2), while the strength of process noise *σ* is determined through the grid search (Supplementary Fig. 2).

#### M step: Optimization of f

The linear filters ***f*** are inferred with the ridge regression where the regularization coefficient is selected as *λ* = 1.0 [38] to avoid overfitting.

Under fixed ***w***^*^, ***f*** is updated as:

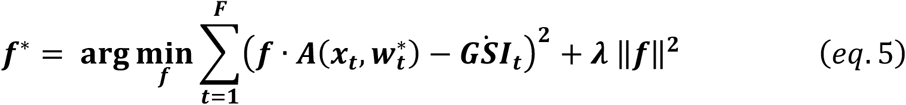

where design matrix ***A***(***x***_***t***_,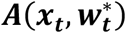) is the set of input signals modulated by ***w***^*^

(Supplementary Note 3). Here we used ***x***_***t***_ to represent [*x*_1,*t*_, *x*_2,*t*_, *x*_3,*t*_, *x*_4,*t*_].

### C. Statistical test

#### The significance test of correlation between the signal importance and gene expression

For estimating the significance of correlation between the signal importances and gene expression patterns, we performed a statistical test with the null hypothesis that the correlation is due to random effects. First, 100,000 randomized trajectories of ***w***_***i***_(= [***w***_***i***,**1**_, ***w***_***i***,**2**_, **…**, ***w***_***i***,***t***_]) were generated while preserving the autocorrelation of the original ***w***_***i***_ using the iterated amplitude adjusted Fourier transformation [39]. The correlation coefficients between the expression pattern of the target gene and each randomized trajectories were then computed, and their histogram was constructed. The null distribution was inferred by applying kernel density estimation. Finally, the p-value was calculated as the probability of obtaining a correlation coefficient greater than the observed value (Supplementary Fig. 4).

## Code availability

ISI is developed under python 3.9.13, and statistical analysis is performed under R 4.4.1. The ISI and the statistical analysis code will be distributed through GitHub after publication.

## Data availability

This study was a re-analysis of existing data (Nakayama et al., PNAS, 2023).

Time-course expression data of medaka were downloaded from the Gene Expression Omnibus (GEO) under accession numbers GSE234401 for the outdoor condition and GSE234565 for the room condition.

https://www.ncbi.nlm.nih.gov/geo/query/acc.cgi?acc=GSE234401.

https://www.ncbi.nlm.nih.gov/geo/query/acc.cgi?acc=GSE234565.

## Author Contributions

H.N., Y.K., T.I., K.A. and T.Y. conceived the project. T.I., H.N. and Y.K. developed the model. A.S., T.N., and T.Y. conducted the experiments. T.I. analyzed the data. T.I., Y.K. and H.N. wrote the manuscript with input from all the authors.

## Acknowledgments

We are grateful to Prof. Nen Saito and all members of the Yoshimura laboratory for the valuable discussions. This study was supported in part by Grant-in-Aid for Transformative Research Areas (B) [grant number 21H05170 to H.N.], Moonshot R&D– MILLENNIA Program [grant number JPMJMS2024-9 to H.N. and Y.K.] by JST, the Cooperative Study Program of Exploratory Research Center on Life and Living Systems (ExCELLS) [program number 21-102 to H.N., 24EXC362, 23EXC323, 21-324, 20-322, 19-318 to T.Y., and 25EXC603 to K.A.], and Japan Society for the Promotion of Science [grant number 23KJ1009 to T.I., and JP25H01362 to K.A.].

