## Supplementary information for "Inverse signal importance in real exposome: How do biological systems dynamically prioritize multiple environmental signals?"

**Supplementary Note. 1** **EM-like algorithm**

The parameters $\boldsymbol{f}$ and $\boldsymbol{w}$**,** in the optimization problem described by *eqs*. 2-5 in the main text, are estimated through an iterative procedure. The procedure approximates the Expectation-Maximization algorithm [25] as detailed below. First, the log likelihood for this model is defined as:

$\begin{aligned} \ln p\left( \mathcal{D} | \boldsymbol{f}, \boldsymbol{w} \right)=\sum_{t=1} \ln\frac{1}{\sqrt{2\pi k^{2}}}\exp\left[ -\frac{\left[ y_{t}(\boldsymbol{x},\boldsymbol{f},\boldsymbol{w})-{\dot{\mathrm{GSI}}}_{t} \right]^{2}}{2k^{2}} \right]\#\left( eq.s1 \right) \end{aligned}$

where $y_{t}(\boldsymbol{x},\boldsymbol{f},\boldsymbol{w})$ is the predicted ${\dot{\mathrm{GSI}}}_{t}$ from our model (eq.2), and $\mathcal{D}$ is a dataset including$\boldsymbol{x}$ and $\dot{\mathbf{GSI}}$.

Then we maximize the log-likelihood $\ln p\left( \boldsymbol{x} | \boldsymbol{f} \right)$, where $\boldsymbol{w}$, latent variables, are marginalized.

***E step:***

In the EM algorithm, the lower bound of the marginalized log-likelihood $\ln p\left( \mathcal{D} | \boldsymbol{f} \right)$ is optimized. In the E step, we compute the lower bound through approximating the posterior $p\left( \boldsymbol{w} | \boldsymbol{x,f} \right)$. Here, for completeness, we will revisit the derivation [25].

$$\ln p\left( \mathcal{D} | \boldsymbol{f} \right)=\ln\sum_{\boldsymbol{w}} p\left( \mathcal{D},\boldsymbol{w} | \boldsymbol{f} \right)$$

$$=\ln\sum_{\boldsymbol{w}} q\left( \boldsymbol{w} \right)\frac{p\left( \mathcal{D},\boldsymbol{w} | \boldsymbol{f} \right)}{q\left( \boldsymbol{w} \right)}$$

$$\begin{aligned} \geq\sum_{\boldsymbol{w}} q\left( \boldsymbol{w} \right)\ln\frac{p\left( \mathcal{D},\boldsymbol{w} | \boldsymbol{f} \right)}{q\left( \boldsymbol{w} \right)} \#\left( eq.s2 \right) \end{aligned}$$

The lower bound follows from Jensen's inequality:

$$\begin{aligned} \ln p\left( \mathcal{D} | \boldsymbol{f} \right)-\sum_{\boldsymbol{w}} q\left( \boldsymbol{w} \right)\ln\frac{p\left( \mathcal{D},\boldsymbol{w} | \boldsymbol{f} \right)}{q\left( \boldsymbol{w} \right)}\geq0 \#\left( eq.s3 \right) \end{aligned}$$

By rearranging the left-hand side, we obtain

$$\ln p\left( \mathcal{D} | \boldsymbol{f} \right)-\sum_{\boldsymbol{w}} q\left( \boldsymbol{w} \right)\ln\frac{p\left( \mathcal{D},\boldsymbol{w} | \boldsymbol{f} \right)}{q\left( \boldsymbol{w} \right)}=\sum_{\boldsymbol{w}} q\left( \boldsymbol{w} \right)\left\{ \ln p\left( \mathcal{D} | \boldsymbol{f} \right)-\ln\frac{p\left( \mathcal{D},\boldsymbol{w} | \boldsymbol{f} \right)}{q\left( \boldsymbol{w} \right)} \right\}$$

$$=\sum_{\boldsymbol{w}} q\left( \boldsymbol{w} \right)\left\{ \ln p\left( \mathcal{D} | \boldsymbol{f} \right)-\ln\frac{p\left( \boldsymbol{w} | \mathcal{D},\boldsymbol{f} \right)p\left( \mathcal{D} | \boldsymbol{f} \right)}{q\left( \boldsymbol{w} \right)} \right\}$$

$$=-\sum_{\boldsymbol{w}} q\left( \boldsymbol{w} \right)\left\{ \ln p\left( \boldsymbol{w} | \mathcal{D},\boldsymbol{f} \right) -\ln q\left( \boldsymbol{w} \right) \right\}$$

$$=\sum_{\boldsymbol{w}} q\left( \boldsymbol{w} \right)\ln\frac{q\left( \boldsymbol{w} \right)}{p\left( \boldsymbol{w} | \mathcal{D},\boldsymbol{f} \right)} (eq. s4)$$

Hence,

$$\begin{aligned} \ln p\left( \mathcal{D} | \boldsymbol{f} \right) -\sum_{\boldsymbol{w}} q\left( \boldsymbol{w} \right)\ln\frac{p\left( \mathcal{D},\boldsymbol{w} | \boldsymbol{f} \right)}{q\left( \boldsymbol{w} \right)}=D_{KL}[q\left( \boldsymbol{w} \right)|\left| p\left( \boldsymbol{w} | \mathcal{D},\boldsymbol{f} \right) \right]\#(eq. s5) \end{aligned}$$

where $D_{KL}$ is Kullback-Leibler divergence.

In the standard EM algorithm, we compute the posterior $q\left( \boldsymbol{w} \right)=p\left( \boldsymbol{w} | \mathcal{D},\boldsymbol{f} \right)$. Then we obtain, since $D_{KL}[q\left( \boldsymbol{w} \right)|\left| p\left( \boldsymbol{w} | \mathcal{D},\boldsymbol{f} \right) \right]=0$, $\ln p\left( \mathcal{D} | \boldsymbol{f} \right)=\sum_{\boldsymbol{w}} q\left( \boldsymbol{w} \right)\ln\frac{p\left( \mathcal{D},\boldsymbol{w} | \boldsymbol{f} \right)}{q\left( \boldsymbol{w} \right)}$.

This means that we can use $\sum_{\boldsymbol{w}} q\left( \boldsymbol{w} \right)\ln\frac{p\left( \mathcal{D},\boldsymbol{w} | \boldsymbol{f} \right)}{q\left( \boldsymbol{w} \right)}$ as the marginalized log-likelihood. Here we adopt the hard EM approximation [28], where we compute the point estimate for $\boldsymbol{w}$ instead of the posterior. We apply Kalman filter & smoother to obtain $\boldsymbol{w}^{*}=\arg\max_{\boldsymbol{w}}\ln p\left( \boldsymbol{w} | \mathcal{D},\boldsymbol{f} \right)$, as detailed in Supplementary Note 2.

***M step:***

Given the hard EM approximation at the E step, we use the mode of complete log-likelihood, $\ln p\left( \mathcal{D},\boldsymbol{w}^{*} | \boldsymbol{f} \right)$, instead of the expectation, $\sum_{\boldsymbol{f}} p\left( \boldsymbol{w} | \mathcal{D},\boldsymbol{f} \right)\ln p\left( \mathcal{D},\boldsymbol{w} | \boldsymbol{f} \right).$ We adopt the ridge regression to solve the following problem,

$\boldsymbol{f}^{*}=\arg\underset{\boldsymbol{f}}{max\{}\ln p\left( \mathcal{D},\boldsymbol{w}^{*} | \boldsymbol{f} \right)+\ln p_{ridge}(\boldsymbol{f})\},$ as detailed in Supplementary Note 3. Then the procedure is iterated.

**Convergence criterion**

The E step and M step is alternatively repeated until the decrease of prediction error falls below ${10}^{-7}$.

**Supplementary Note. 2** **Kalman filter: Algorithm and parameter settings.**

Time-variable coefficient, $\boldsymbol{w}$ is estimated with the Kalman filter for the time discretization of eq. 2 (state-space model).

The state space model is as follows:

$$\left\{ \begin{aligned} \boldsymbol{w}_{t}\boldsymbol{=} \boldsymbol{w}_{t-dt}+\boldsymbol{s} \boldsymbol{s\sim}N\left( \boldsymbol{0,}\sigma dt \boldsymbol{I} \right) \\ \mathbf{y}_{\boldsymbol{t}}=\boldsymbol{H}\left( \boldsymbol{x}_{\boldsymbol{t}},\boldsymbol{f}^{\boldsymbol{*}} \right)\boldsymbol{w}_{\boldsymbol{t}}+\boldsymbol{v} \boldsymbol{v\sim}N\left( \boldsymbol{0, R} \right) \end{aligned} (eq. s6) \right.$$

For obtaining the generalized result, Kalman filter is performed to the three or two data set (see Method: data splitting). In this section, we assume the case of three datasets:

$$\begin{aligned} \boldsymbol{y}_{\boldsymbol{t}}=\left[ \begin{matrix} y_{t}^{\left( \mathrm{Data}1 \right)} & y_{t}^{\left( \mathrm{Data}2 \right)} & y_{t}^{\left( \mathrm{Data}3 \right)} \end{matrix} \right]^{T}\boldsymbol{.} \end{aligned}$$

The observe function is defined as the matrix of convolution of $\boldsymbol{x}_{t}$ and $\boldsymbol{f}^{\boldsymbol{*}}$**:**

$$\begin{aligned} \boldsymbol{H}\left( \boldsymbol{x}_{\boldsymbol{t}},\boldsymbol{f}^{\boldsymbol{*}} \right)=\left[ \begin{aligned} \begin{matrix} \sum_{\tau=0}^{T} f_{1,\tau}^{*}\cdot x_{1,t-\tau}^{\left( Data 1 \right)} & \sum_{\tau=0}^{T} f_{1,\tau}^{*}\cdot x_{1,t-\tau}^{\left( Data 2 \right)} & \sum_{\tau=0}^{T} f_{1,\tau}^{*}\cdot x_{1,t-\tau}^{\left( Data 3 \right)} \\ \sum_{\tau=0}^{T} f_{2,\tau}^{*}\cdot x_{2,t-\tau}^{\left( Data 1 \right)} & \sum_{\tau=0}^{T} f_{2,\tau}^{*}\cdot x_{2,t-\tau}^{\left( Data 2 \right)} & \sum_{\tau=0}^{T} f_{2,\tau}^{*}\cdot x_{2,t-\tau}^{\left( Data 3 \right)} \\ \sum_{\tau=0}^{T} f_{3,\tau}^{*}\cdot x_{3,t-\tau}^{\left( Data 1 \right)} & \sum_{\tau=0}^{T} f_{3,\tau}^{*}\cdot x_{3,t-\tau}^{\left( Data 2 \right)} & \sum_{\tau=0}^{T} f_{3,\tau}^{*}\cdot x_{3,t-\tau}^{\left( Data 3 \right)} \\ \sum_{\tau=0}^{T} f_{4,\tau}^{*}\cdot x_{4,t-\tau}^{\left( Data 1 \right)} & \sum_{\tau=0}^{T} f_{4,\tau}^{*}\cdot x_{4,t-\tau}^{\left( Data 2 \right)} & \sum_{\tau=0}^{T} f_{4,\tau}^{*}\cdot x_{4,t-\tau}^{\left( Data 3 \right)} \end{matrix} \\ \end{aligned} \right]^{T}\boldsymbol{\#}\left( eq. s7 \right) \end{aligned}$$

$\boldsymbol{w}_{\boldsymbol{t}}=\left[ \begin{matrix} w_{1,t} & w_{2,t} & w_{3,t} & w_{4,t} \end{matrix} \right]^{T}$.

Note that while the input signal $x_{i}$ ($i=1$ SR, $i=2$ WT, $i=3$ DL) is common across datasets ($Data 1$, $Data 2$, and $Data 3$), $x_{4}$ (the past GSI) differs among datasets. The process noise $\boldsymbol{s}\sim N\left( \boldsymbol{0},\sigma dt \boldsymbol{I} \right)$ affects both the smoothness of $w^{*}$ and the width of confidential interval for the estimated waves. The observation noise is given by $v\sim N\left( \boldsymbol{0},\boldsymbol{R} \right)$. The optimized time-invariant coefficient $f^{*}$ is an estimated using ridge regression in E step.

In the initial step, we set $w_{i,t}^{*}$ equal to 1 for all $i$ and $t$, then proceed to the M-step.

***Algorithm:***

From the state space model, $w$ evolves as follows with the variance $P$:

$$\begin{aligned} \left\{ \begin{aligned} \boldsymbol{w}_{k+1|k}=\boldsymbol{I}\boldsymbol{w}_{k|k} \\ \boldsymbol{P}_{k+1|k}=\boldsymbol{P}_{k|k}+\sigma\cdot dt\boldsymbol{I} \end{aligned} \right.\#\left( eq. s8 \right) \end{aligned}$$

where $\boldsymbol{w}_{k+1|k}\sim N\left( \boldsymbol{w}_{k+1}, \boldsymbol{P}_{k+1} \right)$.

$\boldsymbol{w}_{k+1|k}$ ($\boldsymbol{P}_{k+1|k}$) is the predicted $\boldsymbol{w}_{k+1}$ ($\boldsymbol{P}_{k+1}$) at step *k,* while $\boldsymbol{w}_{k|k}$ ($\boldsymbol{P}_{k|k}$) is the updated value at step *k*.

The true value of $\boldsymbol{w}$ ($\boldsymbol{P}$) in the step *k*+1 is $\boldsymbol{w}_{k+1} \left( \boldsymbol{P}_{k+1} \right)$. Hence, the predicted value $\boldsymbol{w}_{k+1|k}$ is distributed around the true value $\boldsymbol{w}_{k+1}$ with the variance $\boldsymbol{P}_{k+1}$. At time points where observed data are available,$\boldsymbol{w}$ and $\boldsymbol{P}$ are updated:

$\left\{ \begin{aligned} \boldsymbol{w}_{k+1|k+1}=\boldsymbol{w}_{k+1|k}+\boldsymbol{K}_{k+1}\left( \mathbf{y}_{k}-\boldsymbol{H}_{k}\boldsymbol{w}_{k+1|k} \right) \\ \boldsymbol{P}_{k+1|k+1}=\boldsymbol{P}_{k+1|k}-\boldsymbol{K}_{k+1}\boldsymbol{H}_{k}\boldsymbol{P}_{k+1|k} \\ \boldsymbol{K}_{k+1}= \boldsymbol{P}_{k+1|k}{\boldsymbol{H}_{k}}^{T}\left( \boldsymbol{H}_{k}\boldsymbol{P}_{k+1|k}{\boldsymbol{H}_{k}}^{T}+\boldsymbol{R} \right)^{-1} \end{aligned} \right.$	(*eq*. s9)

The initial state distribution is $\delta$($\boldsymbol{w}$ – $\boldsymbol{w}_{0}$). We determine $\boldsymbol{w}_{0}$ through the following procedure to prevent artificial relaxation of $\boldsymbol{w}$ at the start. First, $\boldsymbol{w}_{0}$ is set to zero. After applying the Kalman filter and the RTS smoother, we update $\boldsymbol{w}_{0}$ to match the final state $\boldsymbol{w}_{\boldsymbol{n|n}}$. With the updated $\boldsymbol{w}_{0}$, we then re-estimate the entire sequence of $\boldsymbol{w}$ using the Kalman filter and RTS smoother. We repeat this update process three times to balance computational costs with convergence.

***Parameter setting:***

***· Observation noise*** $\boldsymbol{v}$

The covariance matrix **R** of the observation noise $\boldsymbol{v}\sim N\left( \boldsymbol{0},\boldsymbol{R} \right)$ is determined based on the properties of the dataset. To this end, first we compute, $\delta_{t}$, the variance of the normalized $\dot{GSI}_{t}$ at time point $t$ among the split datasets. Then the variance is determined as the maximum, $\delta_{max}=max(\delta_{t})$, for numerical stability. Specifically,

***R****=*$\left[ \begin{matrix} \delta_{max} & cov & cov \\ cov & \delta_{max} & cov \\ cov & cov & \delta_{max} \end{matrix} \right]$ where $\delta_{max}=max(\delta_{t})$.

The cov represents the covariance among the three datasets. In our dataset, $\delta_{max}=1.5$ and $cov=0.35$.

***·Process noise*** $\boldsymbol{s}$

Through the comprehensive search, we choose the process noise such that minimizes prediction error.

**Supplementary Note. 3:** **Ridge regression**

Time-invariant coefficient $\boldsymbol{f}$ is estimated using ridge regression.

To avoid overfitting, we set regularization coefficient as $\lambda=1.0$.

The optimal $\boldsymbol{f}$ is estimated by minimizing the residual sum of squares (RSS) while penalizing the sum of squared values of $\boldsymbol{f,}\left\| \boldsymbol{f} \right\|^{\boldsymbol{2}}$**.** For obtaining the generalized result, ridge regression is performed to the three or two data set (see Method: data splitting). In this section, we assume the case of three data set.

$$\begin{aligned} \arg\min\boldsymbol{E}\boldsymbol{=}\sum_{\boldsymbol{t}\boldsymbol{=}\boldsymbol{1}}^{\boldsymbol{F}} \left( \boldsymbol{f}\boldsymbol{\cdot A}\left( \boldsymbol{x}_{t}\boldsymbol{,}\boldsymbol{w}_{t}^{\boldsymbol{*}} \right)\boldsymbol{-}\boldsymbol{y}_{\boldsymbol{t}} \right)^{\boldsymbol{2}}\boldsymbol{+\lambda}\left\| \boldsymbol{f} \right\|^{\boldsymbol{2}}\boldsymbol{\#}\left( eq\boldsymbol{.}s10 \right) \end{aligned}$$

where $\boldsymbol{y}_{\boldsymbol{t}}$ represents temporal differentiation of GSI across three datasets:

$$\begin{aligned} \boldsymbol{y}_{\boldsymbol{t}}=\left[ \begin{matrix} y_{t}^{\left( \mathrm{Data}1 \right)} & y_{t}^{\left( \mathrm{Data}2 \right)} & y_{t}^{\left( \mathrm{Data}3 \right)} \end{matrix} \right] \end{aligned}$$

The regression parameter $\boldsymbol{f}$ is defined as:

$$\begin{aligned} \boldsymbol{f}\boldsymbol{=}\left[ \begin{matrix} \left\{ f_{1,t} \right\}_{t=0}^{t=T} & \left\{ f_{2,t} \right\}_{t=0}^{t=T} & \left\{ f_{3,t} \right\}_{t=0}^{t=T} & \left\{ f_{4,t} \right\}_{t=0}^{t=T} \end{matrix} \right]\boldsymbol{\#} \end{aligned}$$

where ${\{f_{i,t}\}}_{t=0}^{t=T}\boldsymbol{=}\left[ f_{i,0}, \ldots,f_{i,T} \right]$**.**

The design matrix $\boldsymbol{A}\left( \boldsymbol{x}_{t}\boldsymbol{,}\boldsymbol{w}_{t}^{\boldsymbol{*}} \right)$ is an expanded form of the multiplication of $\boldsymbol{x}_{t}$ and $\boldsymbol{w}_{t}^{\boldsymbol{*}}$:

$$\begin{aligned} \boldsymbol{A}\left( \boldsymbol{x}_{t}\boldsymbol{,}\boldsymbol{w}_{t}^{\boldsymbol{*}} \right)=\left[ \begin{aligned} \begin{matrix} w_{1,t}^{*}\left\{ x_{1,t-\tau}^{\left( Data 1 \right)} \right\}_{\tau\boldsymbol{=}0}^{\tau=T} & w_{1,t}^{*}\left\{ x_{1,t-\tau}^{\left( Data 2 \right)} \right\}_{\tau\boldsymbol{=}0}^{\tau=T} & w_{1,t}^{*}\left\{ x_{1,t-\tau}^{\left( Data 3 \right)} \right\}_{\tau\boldsymbol{=}0}^{\tau=T} \\ w_{2,t}^{*}\left\{ x_{2,t-\tau}^{\left( Data 1 \right)} \right\}_{\tau\boldsymbol{=}0}^{\tau=T} & w_{2,t}^{*}\left\{ x_{2,t-\tau}^{\left( Data 2 \right)} \right\}_{\tau\boldsymbol{=}0}^{\tau=T} & w_{2,t}^{*}\left\{ x_{2,t-\tau}^{\left( Data 3 \right)} \right\}_{\tau\boldsymbol{=}0}^{\tau=T} \\ w_{3,t}^{*}\left\{ x_{3,t-\tau}^{\left( Data 1 \right)} \right\}_{\tau\boldsymbol{=}0}^{\tau=T} & w_{3,t}^{*}\left\{ x_{3,t-\tau}^{\left( Data 2 \right)} \right\}_{\tau\boldsymbol{=}0}^{\tau=T} & w_{3,t}^{*}\left\{ x_{3,t-\tau}^{\left( Data 3 \right)} \right\}_{\tau\boldsymbol{=}0}^{\tau=T} \\ w_{4,t}^{*}\left\{ x_{4,t-\tau}^{\left( Data 1 \right)} \right\}_{\tau\boldsymbol{=}0}^{\tau=T} & w_{4,t}^{*}\left\{ x_{4,t-\tau}^{\left( Data 2 \right)} \right\}_{\tau\boldsymbol{=}0}^{\tau=T} & w_{4,t}^{*}\left\{ x_{4,t-\tau}^{\left( Data 3 \right)} \right\}_{\tau\boldsymbol{=}0}^{\tau=T} \end{matrix} \\ \end{aligned} \right]\boldsymbol{\#}\left( eq. s11 \right) \end{aligned}$$

where $w_{i,t}^{*}{\boldsymbol{\{}x_{i,t-\tau}\boldsymbol{\}}}_{\tau\boldsymbol{=}0}^{\tau=T}\boldsymbol{=}\left[ w_{i,t}^{*} x_{i,t}, \ldots,w_{i,t}^{*} x_{i,t-T} \right]^{T}$**.**

Note that while the input signal $x_{i}$ ($i=1$ SR, $i=2$ WT, $i=3$ DL) is common across datasets ($Data 1$, $Data 2$, and $Data 3$), $x_{4}$ (the past GSI) differs among datasets. The weight $w_{i,t}^{*}$ is a time-variant coefficient estimated through the Kalman filter in E step. In the initial step, $w_{i,t}^{*}=1$ for all $i$ and $t$.

**Supplementary Figures**

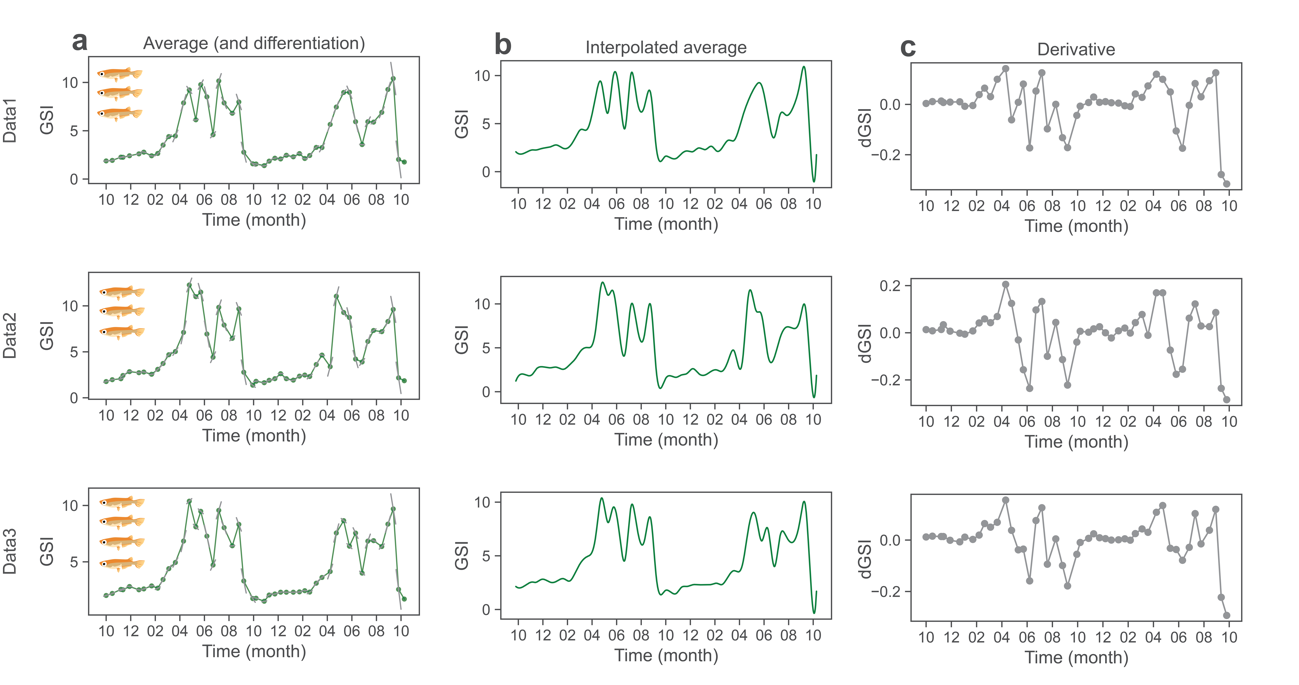

**Supplementary Figure. 1** **Data splitting and processing.** The 10-sample dataset is split into Data1 (n=3, upper row), Data2 (n=3, middle row), and Data3 (n=4, lower row). **a** Average GSI in each dataset. The differentiation around each sampling point is denoted with a grey line. **b** Interpolated GSI. Average GSI is converted to the 1-day interval. **c)** Derivative of GSI. Y-axis is the slope of grey line in **a.**

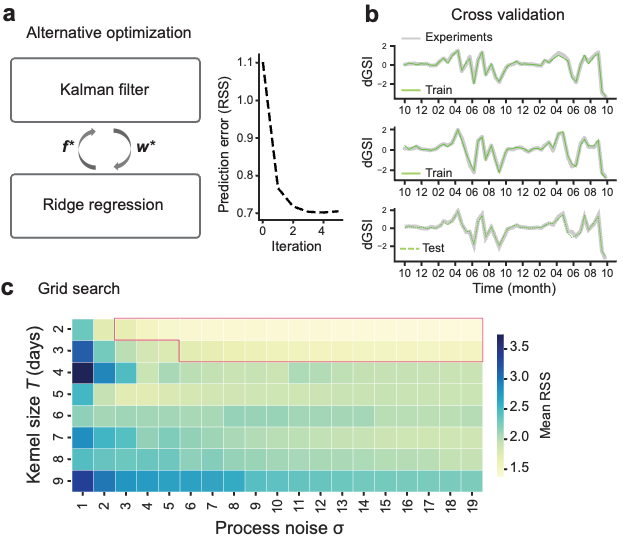

**Supplementary Figure 2. Parameter Optimization**. **a** **Alternative optimization:** The prediction error decreases through iterations (right). **b** **Cross-Validation:** Two datasets are used for train and the remaining data is used for test to prevent overfitting of hyperparameters. The fitting results for train (upper and middle) and test (lower) under the best hyperparameters are shown. **c** **Grid Search:** Kernel size *T* and the strength of process noise $\sigma$ are optimized by evaluating the mean residual sum of squares (RSS) across all cross-validation sets. Cells with the lowest RSS values (rounded to one decimal place) are highlighted in red. Among these, those with kernel size greater than 2 are averaged to derive a representative waveform that captures meaningful structure in the response function.

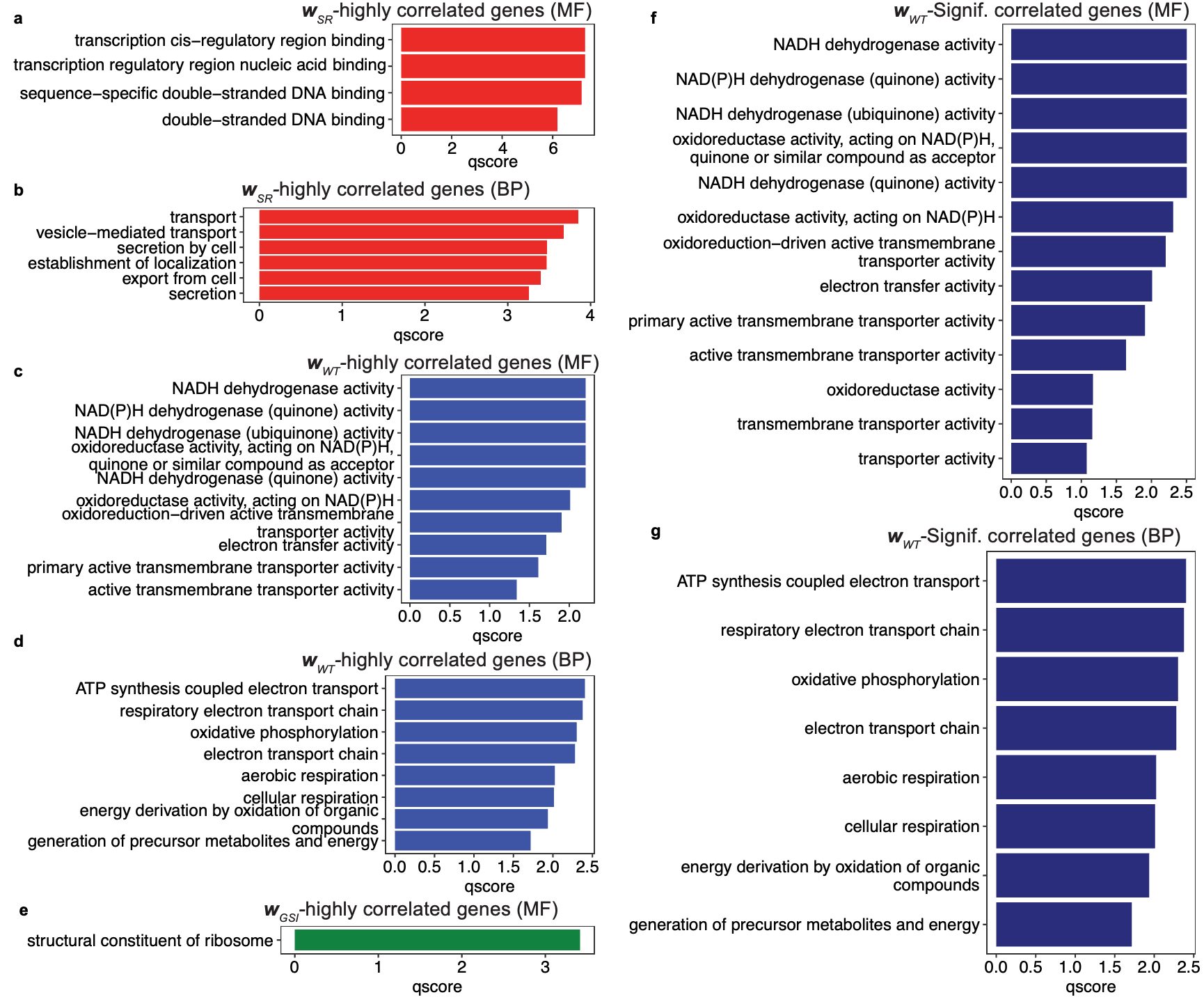

**Supplementary Figure 3. GO enrichment analysis**. The horizontal axis represents the significance of enrichment qscore ( -log_10_p ), while the vertical axis indicates the enriched functions (q > 1). **a,b,c**,**d** and **e**) GO enrichment analysis of biological process (BP) or molecular function (MF) for genes highly correlated with $\boldsymbol{w}_{SR}$(**a**: MF, **b**: BP)), $\boldsymbol{w}_{WT}$ (**c**: MF, **d**: BP), and $\boldsymbol{w}_{GSI}$ (**e**: MF). **f** and **g**) GO enrichment analysis for genes significantly correlated with $\boldsymbol{w}_{WT}$ (**f** MF, **g**: BP).

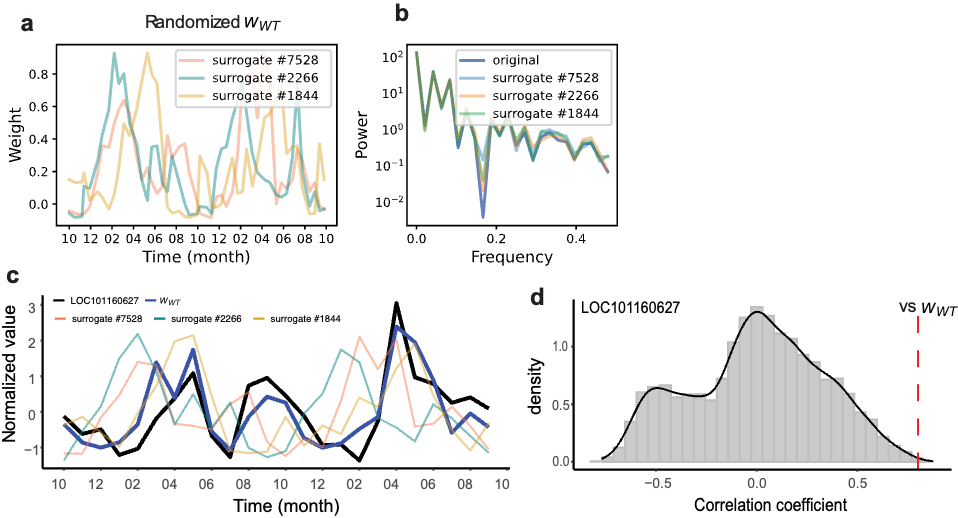

**Supplementary Figure 4. Statistical test for the correlation of gene expression pattern and signal importance. a** Randomized signal importance. Three examples of randomized $\boldsymbol{w}_{WT}$ are shown. **b** Power spectra of the randomized waveforms in (a). The overlap among the spectra confirms the preservation of autocorrelation. **c** Comparison of the gene expression pattern (LOC101160627; black), signal importance $\boldsymbol{w}_{WT}$ (blue), and randomized waveforms (the other colors). **d** The distribution of correlation coefficients between expression pattern of LOC101160627 and each randomized $\boldsymbol{w}_{WT}$ (n=100,000). The estimated distribution function using kernel density estimation is denoted with black line. The correlation coefficient with the original waveform ($\boldsymbol{w}_{WT}$) is indicated with the red dashed line. The statistical significance is quantified as the upper-tail probability beyond the red line.

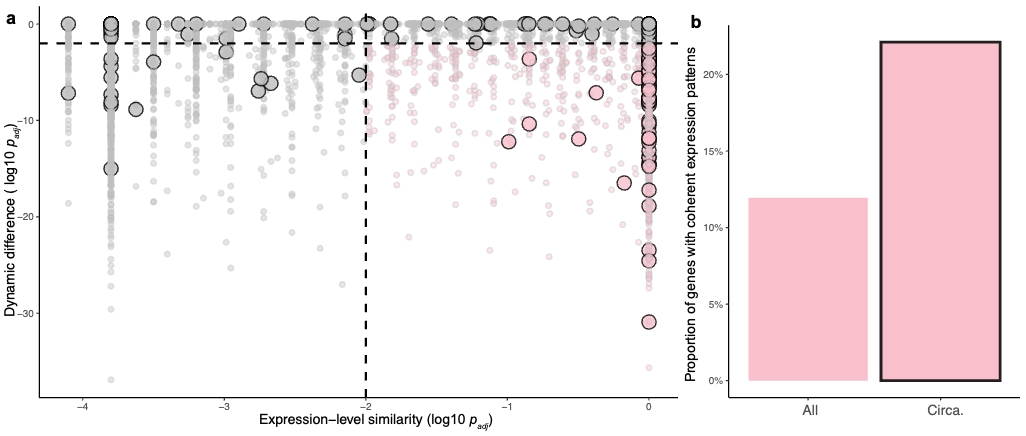

**Supplementary Figure 5. Circannual genes tend to exhibit coherent expression dynamics between outdoor and indoor conditions.** **a** Comparison between expression level similarity and temporal pattern differences across conditions (room and outdoor). The x-axis shows the adjusted p-values (log₁₀ scale) from Wilcoxon tests assessing similarity in expression levels; the y-axis shows adjusted p-values from Spearman tests evaluating dynamic difference. Each point represents a gene, and larger circles indicate circannual genes identified in the previous study (Nakayama et al., 2023). Genes located in the lower right quadrant (highlighted in pink) are considered to exhibit coherent expression dynamics across the two conditions. **b** Proportion of genes with coherent expression profiles among all genes (left) and circannual genes (right). Note that the limited number of genes below the threshold is an expected consequence given that conservative statistical test. A Fisher’s exact test indicates a significant enrichment of coherence among circannual genes (p = 4.47 × 10⁻^8^).

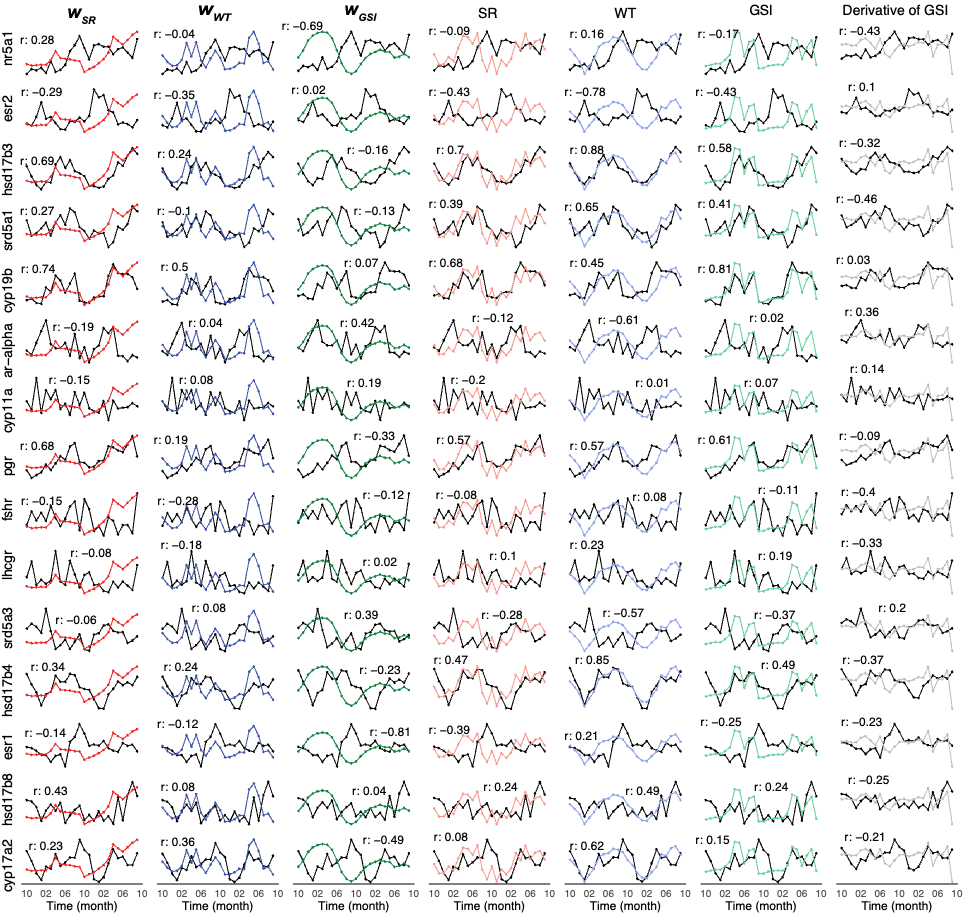

### **Supplementary Figure 6.** **Covariation patterns between sex-hormone transcripts, environmental signals, and inferred signal importance weights.** Row represents a transcript involved in sex hormone synthesis, signaling, or regulation (n = 15; see Supplementary Table 2). Columns show the inferred signal importance weights ($\boldsymbol{w}_{SR}$, $\boldsymbol{w}_{WT}$, $\boldsymbol{w}_{GSI}$) and raw environmental variables (SR, WT, GSI, ΔGSI) across monthly time points. In each panel, the horizontal axis indicates time (months; two years), and the vertical axis indicates the normalized value (FPKM or weight). The Pearson correlation coefficient (r) between each gene and variable is annotated beside each panel.

**Supplementary Table 1. Gene lists showing high correlation with each signal importance.**

Genes that exhibit high correlation (>0.6) with each signal importance ($w_{SR}$, $w_{WT}$, $w_{DL}$, and $w_{GSI}$) are listed. Separate sheets are provided for each signal importance. The first and second columns show the gene symbol and gene ID, respectively. The third column reports the correlation coefficient, followed by the p-value and the adjusted p-value (Bonferroni correction). Note that p-values were estimated as described in Supplementary Figure 4.

**Supplementary Table 2.** **Gene list involved in sex hormone.**

Manually curated transcripts related to sex hormones are listed. The first and second columns show the gene symbol and gene ID, respectively. The third column reports the function based on KEGG orthology, and the fourth column reports the function based on the NCBI NR database description. The fifth column provides literature evidence for involvement in sex hormone function.
